# Fluid-structure interaction simulation of tissue degradation and its effects on intra-aneurysm hemodynamics

**DOI:** 10.1101/2021.09.01.458529

**Authors:** Haifeng Wang, Klemens Uhlmann, Vijay Vedula, Daniel Balzani, Fathollah Varnik

**Affiliations:** Interdisciplinary Centre for Advanced Materials Simulation (ICAMS), Ruhr-Universität Bochum, 44801 Bochum, Germany; Continuum Mechanics, Ruhr-Universität Bochum, 44801 Bochum, Germany; Department of Mechanical Engineering, Columbia University in the City of New York, 10027 New York, United States

## Abstract

Tissue degradation plays a crucial role in vascular diseases such as atherosclerosis and aneurysms. Computational modeling of vascular hemodynamics incorporating both arterial wall mechanics and tissue degradation has been a challenging task. In this study, we propose a novel finite element method-based approach to model the microscopic degradation of arterial walls and its interaction with blood flow. The model is applied to study the combined effects of pulsatile flow and tissue degradation on the deformation and intra-aneurysm hemodynamics. Our computational analysis reveals that tissue degradation leads to a weakening of the aneurysmal wall, which manifests itself in a larger deformation and a smaller von Mises stress. Moreover, simulation results for different heart rates, blood pressures and aneurysm geometries indicate consistently that, upon tissue degradation, wall shear stress increases near the flow-impingement region and decreases away from it. These findings are discussed in the context of recent reports regarding the role of both high and low wall shear stress for the progression and rupture of aneurysms.

## 1 Introduction

Microscopic degradation in vascular tissue is a severe pathological process and is associated with vascular diseases such as aneurysms with high mortality [1–3].

Hemodynamic forces, such as pressure and wall shear stress (WSS, defined as the tangential stress component induced by the flowing blood on the vascular walls), play a crucial role in vascular physiology and pathology by affecting the biochemical response of the surrounding tissue [4–6]. Arterial wall tissue gradually changes under abnormal WSS and other biological factors (long-term effect). The resulting change of arterial geometry, on the other hand, alters the flow characteristics (short-term effect). Both low WSS [7–9] and high WSS [10–12] have been associated with the progression and rupture of aneurysms. To reconcile these conflicting observations, Meng et al. [13] have proposed two independent pathways for the mechanistic link between hemodynamic forces and aneurysm’s pathological changes. Within this hypothesis, low WSS is related to the inflammatory cell-mediated destructive remodeling of the arterial wall; high WSS, which is prevalent near the flow impingement site, is associated with the mural cell-mediated remodeling [13]. Mural cells, including smooth muscle cells and fibroblasts, are essential components of blood vessels. High WSS can trigger biochemical responses leading to the production and activation of proteases, in particular, matrix metalloproteinases production through mural cells [13]. High blood pressure is also known to be responsible for degradation and enlargement of the aneurysm.

Despite their significance, however, it is difficult to clinically measure the spatial distribution and temporal evolution of WSS and pressure inside the aneurysm dome. Numerical simulations provide a promising alternative. In a recent study [14] we have shown that blood flow inside an aneurysm exhibits complex features (such as flow disturbances) associated with wall softness. Also fluid-structure interaction (FSI) simulations with complex wall models [15] have unveiled significant effects on WSS even in simplified arteries. To date, however, we are not aware of any numerical study that couples hemodynamics with the progression of tissue degradation and vessel wall softening.

We propose a novel FSI model capable of accounting for the interaction of blood flow with arterial walls and simultaneously capturing the degradation of vascular tissue. The coupled model is first benchmarked on a canonical problem and is then used to study the effect of tissue degradation on the spatial distribution of strain, WSS and pressure in aneurysms. Upon degradation, we observe a weakening of the vessel wall resulting in larger strain and smaller stress magnitude. Remarkably, WSS increases (up to 20%) near the flow-impingement region but decreases away from it. Near-wall pressure, however, changes only slightly upon tissue degradation.

## 2 Methods

We have developed a new computational framework by combining a tissue degradation model [16, 17] and a finite element-based FSI solver [18]. A brief description of the individual components is given below.

### 2.1 Tissue degradation model

For the modeling of the arterial tissue, the degradation model [16] is used and briefly recapitulated here. The model is formulated in terms of an extended continuum damage mechanics approach, where a strain energy density Ψ is defined as a function of the right Cauchy-Green deformation tensor ***C*** = ***F***^T^***F***, with ***F*** denoting the deformation gradient tensor. Based thereon, the Cauchy stress tensor ***σ*** can be computed from the second Piola-Kirchhoff stress tensor ***S*** = 2*∂*_***C***_Ψ by ***σ*** = (det***F***)^*−*1^***FSF***^*T*^. The total strain energy function is chosen to be

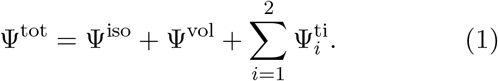

The first term on the rhs of Eq. (1) is the isotropic part describing the ground matrix material, also known as elastin, which is modeled by the incompressible Neo-Hookean model 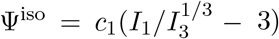. The second term is the compressible function 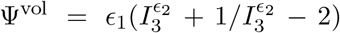 [19], serving here as volumetric penalty function to account for near-incompressibility. Here, *I*_1_ = tr***C*** and *I*_3_ = det***C*** denote the first and the third invariants of the right Cauchy-Green tensor ***C***. The effect of damage is accounted for in the last part of Eq. (1) namely, the transversely isotropic part, which describes the material behavior of the collagen fibers in the tissue. A so-called damage function *D*_*i*_ is introduced via

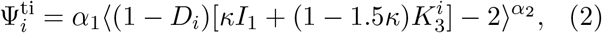

with *i* = 1, 2 indexing fiber families (i.e., each fiber family, *i*, has its own damage parameter, *D*_*i*_). The Macaulay brackets, ⟨(·)⟩ = [(·) + |(·)|] /2, filter out positive values. Here, 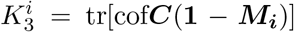 is the fundamental polyconvex function, introduced by Schröder and Neff [20], with the definition of the structural tensor ***M***_***i***_ = ***A***_***i***_ ⊗ ***A***_***i***_, given in terms of each fiber direction vector ***A***_*i*_. The cofactor is defined as cof***C*** = det***CC***^*−*1^. In the case where no damage occurs, i.e. *D*_*i*_ = 0, the energy function corresponds to the one proposed in [21] which is extended by the fiber dispersion approach from [22]. The energy is polyconvex and it a *priori* ensures material stability and thus, a physically reasonable material behavior [23]. Here, *c*_1_, *∈*_1_, *∈*_2_, *α*_1_, *α*_2_ and *κ* are material parameters. *D*_*i*_ are scalar damage functions for each of the mainly two fiber families *i* = 1, 2.

The scalar damage functions, *D*_*i*_, for each fiber family *i* = 1, 2, used to capture remnant strains (i.e., strain at zero stress level after unloading) within the fibers and the *stress-softening* effect, is defined as

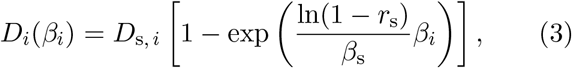

where the maximally reachable damage value for fixated load levels is denoted by *D*_s,*i*_ ∈ [0, 1). The fraction of the maximum damage *r*_s_ is set to *r*_s_ = 0.99, as in [16]. *β*_s_ *>* 0 is the value of the internal variable *β*_*i*_ corresponding to damage saturation. The internal variable is defined as 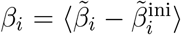, With 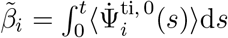 allowing for continuous damage evolution for loading and re-loading paths and 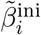 denoting the damage initiation threshold. The latter is set equal to 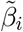 in the scenario where the transition from physiological to supra-physiological loading is reached. Thereby, it is a locally distributed quantity depending on the boundary value problem under investigation and not a material parameter. *t* indicates the time at the current state, and 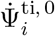 is the first time derivative of the fictitiously undamaged (effective) strain energy density 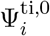 in fiber direction *i*. The maximally reachable damage value for fixated load levels *D*_s, *i*_ is expressed as

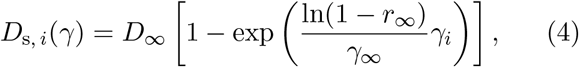

with the predefined converging limit for the overall damage value *D*_*∞*_ ∈ [0, 1) and *γ*_*∞*_ *>* 0 representing the value of the internal variable *γ*_*i*_ reached at the limit fraction *r*_*∞*_ = 0.99. Note that *r*_*s*_ as well as *r*_*∞*_ represent the fraction of the maximal damage values *D*_*s,i*_, i.e. *D*_*∞*_, at which the internal variables *β*_*i*_ and *γ*_*i*_, respectively, reach the values of the material parameters *β*_*s*_, i.e. *γ*_*∞*_. Therefore, *r*_*s*_ as well as *r*_*∞*_ should not be considered as material parameters, but rather as parameters defining the physical interpretation of the material parameters *β*_*s*_ and *γ*_*∞*_.

In order to ensure that *D*_s, *i*_ remains unchanged during a cyclic process under fixed maximum load levels, the second internal variable

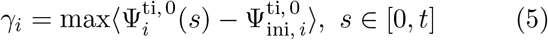

is defined as the maximum value of the effective energy reached up to the current state at *t*. 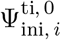 denotes the effective strain energy density at an initial damage state obtained at the limit of the physiological domain. The damage saturation criterion is expressed as

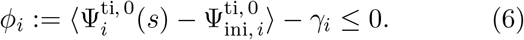

Various alternative approaches for the description of tissue degradation have been proposed in the literature, see e.g. [24–27]. The degradation model considered here has been validated in previous studies [16, 17] and proved to correspond well with cyclic experiments performed on different types of arteries. An application of the model to aneurysms is, however, new. In this context, it is noteworthy that pathological tissue remodeling often plays a key role in cerebral aneurysms. The model used in the present study does not account for this effect. However, the focus of the present investigation is not on long-time degradation processes, where tissue remodeling is of key importance, but on the combined effect of flow-induced forces and short-time degradation which occurs during a few cardiac cycles. Thus, this study has a qualitative character and presents a first step in a long-time effort towards a more realistic model. One of the advantages of such numerical investigations is that they allow to identify potentially important issues and thus reduce the number and extent of clinical studies to a small set of more promising scenarios. The present study emphasizes via a set of extensive simulations the importance of coupling blood flow with tissue degradation and hopes to provide guidance to future clinical studies.

In numerical simulations, a loss of ellipticity of the governing differential equations may be observed when including material stiffness degradation, which results in mesh-dependent simulations. In this case, either a relaxed damage formulation [28] or a gradient-enhanced approach [29] may be applied. In the present paper, however, numerical analysis has shown mesh-convergence and thus, such more advanced formulations were not considered.

### 2.2 FSI solver

We approximate blood as an incompressible Newtonian fluid. For fluid-structure interactions, we employ the well-developed, validated and open-sourced finite element method-based solver, Sim-Vascular/svFSI [18, 30], which uses an arbitrary Lagrangian-Eulerian formulation of Navier-Stokes equations to model incompressible flows on moving domains [31]. It has been applied to a variety of applications including blood flow in developing ventricles [5], patient-specific studies including pulmonary hypertension and coronary artery bypass graft [32], and FSI simulations in aortic dissections [33]. For further details on the methodology and implementation, the reader is referred to [31, 33].

For the flow boundary conditions, we prescribe a pulsatile flow at the inlet with a parabolic velocity profile (Fig. 1). We prescribe a three-element Windkessel-type boundary condition at each outlet and the parameters are tuned to match the systolic, diastolic and mean blood pressures (subsection 2.4). At each cross-sectional end of the structural domain, we apply homogeneous Dirichlet boundary conditions to anchor its location.

**Fig 1:**
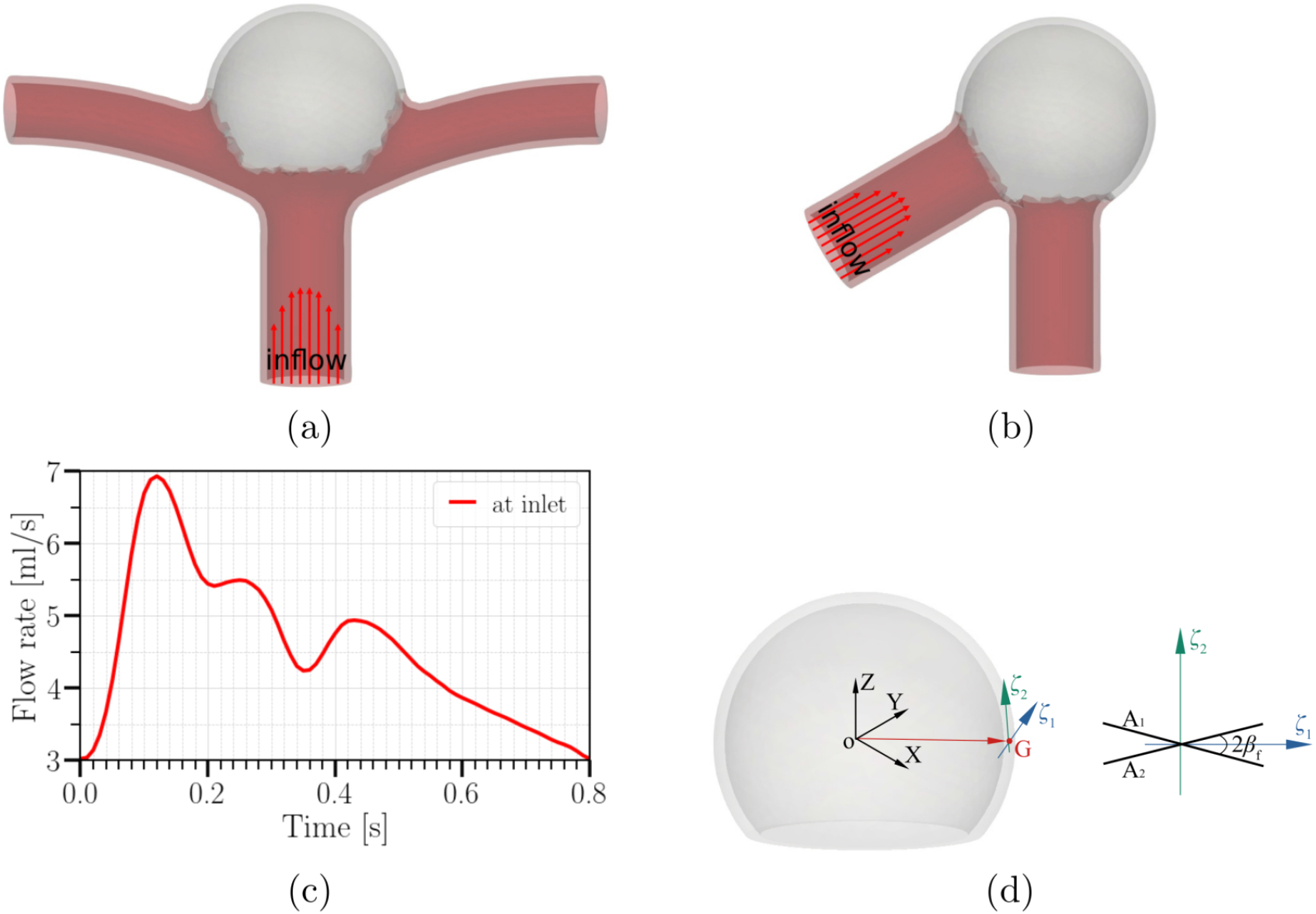
(a–b) Geometries studied in this work; (a) an idealized internal carotid artery bifurcation aneurysm (based on the data in [34, 35]); (b) idealized aneurysm based on the data in [36]. The spherical caps (aneurysms) in (a) and (b) have identical sizes; the diameter of the two vessels in (b) is the same as that of the parent vessel in (a). Arrows highlight the parabolic velocity profile imposed at the inlet. The tissue degradation model is applied only on the aneurysmal region (colored in grey); the remaining of the vessel (red) is modeled as Neo-Hookean. (c) Temporal variation of the imposed flow rate at the inlet [37]. (d) Fiber orientations in a spherical aneurysm (as adopted by [38]). Two fiber families, ***A***_1_, ***A***_2_, are applied at each Gauss integration point G, where 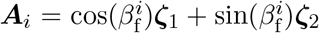 with *i* = 1, 2 and 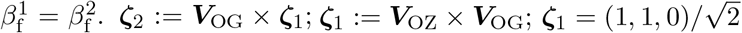 is used in the case that ***V***_OG_ is parallel to ***V***_OZ_.

### 2.3 Verification of the coupling between the damage model and the FSI-solver

The coupling between our in-house tissue damage subroutine and the FSI is facilitated via the exchange of the strain and stress fields. The deformation gradient tensor, ***F*** of the vessel wall computed at the integration point-level and other userprovided parameters are passed as input to our inhouse material subroutine. The subroutine then returns the second Piola Kirchhoff stress tensor and the corresponding elasticity tensors to the FSI solver. The FSI-solver Simvascular/svFSI, in turn, solves for the displacement field of the vessel wall and the blood velocities and pressure, using a monolithic coupling between the structural dynamics and the Navier-Stokes equations.

To validate the coupling between the FSI solver and the tissue-degradation material routine, we use a canonical problem where a cube is subjected to a uniaxial tension (see Fig. 2(a)). The Cauchy stress versus strain profiles, without and with the damage model, obtained using the proposed numerical framework are compared against those obtained from FEAP software package [16]. However, as FEAP is primarily a solid mechanics software, we do not model the blood flow for this verification exercise. Instead, we use svFSI as a pure solid mechanics solver that interfaces with our in-house material subroutine. All relevant simulation parameters are given in Table 1.

**Table 1:** Material parameters for a human carotid artery [16]. For the parent and branch vessels outside the aneurysm (Fig. 1(a-b)), elastic modulus and Poisson’s ratio are chosen as 1000 kPa [39] and 0.49, respectively.

| $c_1$ [kPa] | $\epsilon_1$ [kPa] | $\epsilon_2$ [-] | $\alpha_1$ [kPa] | $\alpha_2$ [-] |
| --- | --- | --- | --- | --- |
| 9.02 | 499.8 | 2.4 | 1400 | 2.2 |
| $\kappa$ [-] | $\beta_f$ [°] | $D_\infty$ [kPa] | $\gamma_\infty$ [kPa] | $\beta_s$ [-] |
| 1e-8 | 39.87 | 0.96 | 17.98 | 0.06 |

**Fig 2:**
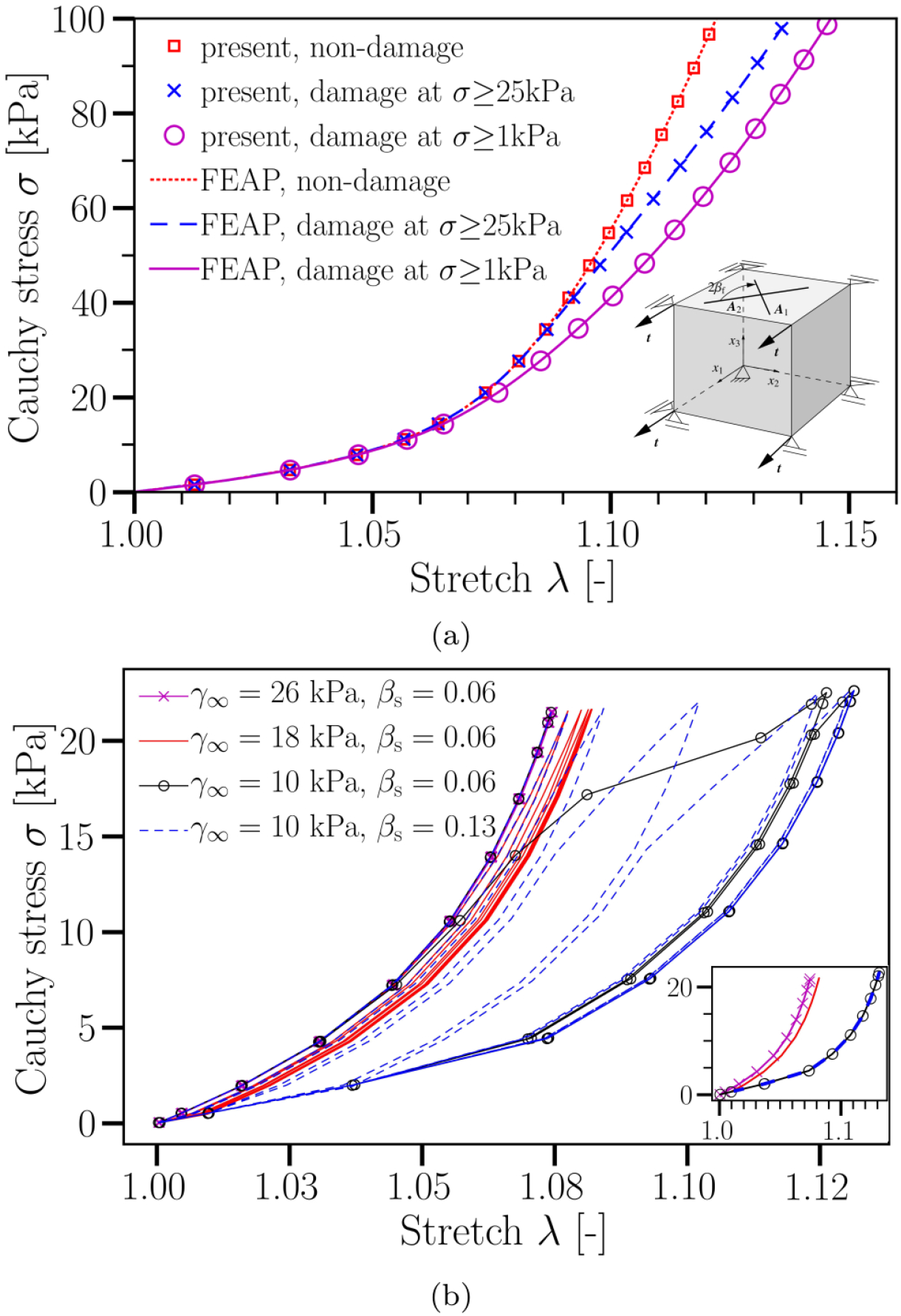
(a) Uniaxial tension tests using a sample with cubic shape without and with damage. Results are shown for the present hybrid model and FEAP. In all cases, the input load (or transmural pressure) is increased from 0 to 100 kPa linearly with time. As can be seen, simulation results using the hybrid FSI framework and FEAP are identical verifying our implementation. All relevant material parameters are given in Table 1 as all cases considered here are quasi-static. Fiber directions are indicated in the inset; the angle between the two fibers, ***A***_*i*_, is 2*β*_f_ with *β*_f_ = 39.87. (b) Cyclic uniaxial tension tests using the same geometry and material parameters as in (a) but different values of *γ*_*∞*_ and *β*_*s*_. The inset in (b) shows the same data as in the main image but for the last cycle only. It highlights that the behavior after many cycles does not depend on *β*_*s*_ but only on *γ*_*∞*_ (note the overlap of data for *β*_*s*_ = 0.06 and *β*_*s*_ = 0.13 in the case of *γ*_*∞*_ = 10 kPa). Each simulation is performed for ten cycles. In all the cases shown, a *cyclic* load is imposed, which varies between 0 and 20 kPa. The degradation process is turned on after the first cycle. As can be seen, the damage saturates after a few (up to a maximum of five) cycles in all the cases investigated; after this initial degradation stage, no further morphological changes occur with time within the present material model. Importantly, the damage parameters, *γ*_*∞*_ and *β*_*s*_, control the degradation intensity and rate, respectively.

We show a very strong agreement in Cauchy stress versus strain profiles between the present numerical framework and FEAP (Fig. 2(a)).

The material parameters used in this study are listed in Table 1. As illustrated in Fig. 2(b), other choices of degradation intensity and the rate of damage saturation can be also simulated by adapting the damage parameters *γ*_*∞*_ and *β*_*s*_, respectively. Indeed, the material parameters given in Table 1 can be easily tuned for long-term tissue degradation, provided that reliable experimental and/or clinical data become available.

### 2.4 Parameters for FSI simulation including material damage

To gain a qualitative understanding of the effects arising from tissue degradation, we perform two sets of simulations: One with degradation switched on and the other switching it off (i.e., assuming a non-degrading tissue), using a bifurcation aneurysm model (Fig. 1(a)).

The inner diameters of the parent vessel and its two branches are 0.41 cm and 0.29 cm [35], respectively. We assume identical branch vessel dimensions to simplify the tuning of the Windkessel parameters (i.e., distal resistance *R*_d_, proximal resistance *R*_p_ and capacitance *C* (RCR)) at each outflow branch. The inner diameter of the spherical cap (aneurysm) is 1.0 cm in this work. Wall thickness is 0.04 cm and spatially homogeneous. All meshes use quadratic tetrahedrons and have a mean effective spatial resolution of 0.07 mm. For each case investigated in the present study, the fluid and solid domains consist of approximately 128,000 nodes / 85,000 tetrahedral elements. The discretized time step is set to 10^*−*4^ s; for a cycle with a duration of 0.8 s, the temporal resolution is 8,000 time-steps per cardiac cycle. Typically, each cardiac cycle runs for approximately 60 hours on 38 cores on a multicore workstation with Intel(R) Xeon(R) Gold 6148 CPU (2.40 GHz).

The time dependence of the flow rate at the inlet is based on the data from [37] (Fig. 1(c)). The mean inlet flow rate is *Q*_in_ = 4.6 ml/s (range: 3.0– 6.9 ml/s). The duration of cardiac cycle is 0.8 seconds. The flow rate at each branch is *Q*_b_ = 0.5*Q*_in_.

Pulse pressure, i.e., the difference between systolic and diastolic blood pressures, in most people is between 40 and 60 mmHg. A small increase (e.g., by 10 mmHg) in the pulse pressure often significantly increases the cardiovascular risk (e.g., by 20%) [40]. In this study, the peak systolic and end diastolic blood pressures are 140 mmHg and 70 mmHg (mean: 100 mmHg), respectively, similar to the hypertension cases reported in [41]. The total resistance at each branch is *R*_tot,b_ = 5.7·10^4^ g cm^*−*4^ s^*−*1^ to meet the pressure shown. The distal and proximal resistance at each outlet is given by *R*_d,b_ = *k*_d_*R*_tot,b_, and *R*_p,b_ = (1−*k*_d_)*R*_tot,b_, where the factor *k*_d_ defines the ratio of distal to total resistance and is fixed for all outlets to *k*_d_ = 0.9 [42]. The capacitance at each branch is identical and equal to 10^*−*6^ cm^4^ s^2^ g^*−*1^. Fluid density and viscosity are *ρ*_f_ = 1.055 g cm^*−*3^ and *η*_f_ = 0.04 g cm^*−*1^ s^*−*1^, respectively. In all FSI simulations, the solid density is *ρ*_s_ = 1.2 g cm^*−*3^.

The fiber orientations applied for the aneurysm dome are indicated in Fig. 1(d). Due to the lack of experimental and clinical data on the fiber orientations of any aneurysms, we set *β*_f_ = 40° as in [16]. The other parameters of the material model are listed in Table 1.

### 2.5 Metrics for analysis

Hemodynamic quantities play an important part in the mechanobiological development of vascular diseases [43]. Among these, WSS and time-averaged wall shear stress (TAWSS) are often reported to be closely associated with the formation and rupture of aneurysms [13, 44].

The TAWSS at an arbitrary position *x* is simply the average magnitude of the WSS vector ***τ***_w_ over one cardiac cycle of duration *T*

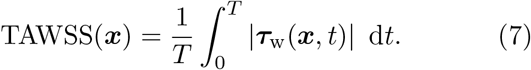

To quantify the role of tissue degradation in aneurysmal wall and intra-aneurysm hemodynamics, we also examine the near-wall pressure, von Mises stress and strain magnitude on the aneurysmal wall. We compute the time average of each metric, defined along similar lines as Eq. 7. We further analyze the relative changes in these metrics due to tissue degradation. Given a time-averaged quantity *f*(*x*) at point *x*, the *relative percentage change* in *f*(*x*) between the degraded (*f*^d^) and non-degraded (*f*^nd^) cases is defined via

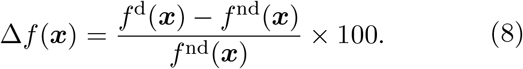

## 3 Results and discussion

In this section, we use the proposed numerical model to investigate the effect of tissue degradation on the hemodynamics and the mechanical response of a bifurcation aneurysm model (Fig. 1(a)). We focus on a simplified aneurysm model to assess the feasibility of performing a FSI simulation combined with a material model and develop preliminary insights into the role of hemodynamics on tissue degradation and vice versa. However, extending our numerical framework to performing patient-specific FSI simulations of tissue degradation is straightforward, provided that personalized data on tissue fiber architecture and damage model parameters become available [45, 46].

The development of an aneurysm usually takes months or even years [47]. This slow process depends on several physiological and hemodynamic parameters, and is far from being completely understood. However, there is strong clinical evidence supporting the hypothesis that local WSS and blood pressure play a critical role in aneurysm development and rupture [13]. At the same time, the lack of sufficient experimental and clinical data has limited the development of models that adequately account for the long-term dynamics of tissue degradation. Moreover, the computational cost of FSI simulations prohibits us from simulating a long-term hemodynamic response to tissue damage, and vice versa. Therefore, we focus on understanding the hemodynamic implications of tissue degradation within a few cardiac cycles.

We ignore the initial transients and focus on the dynamics after five cardiac cycles.

### 3.1 Effects of tissue degradation

Figure 3 shows the spatial distribution of WSS and hydrostatic pressure at two instants (peak systole and end diastole) both in the presence and absence of tissue degradation. A close scrutiny of these images provides some first hints towards the effect of tissue degradation but it seems at this stage a bit too early to draw any specific conclusion based on these images.

**Fig 3:**
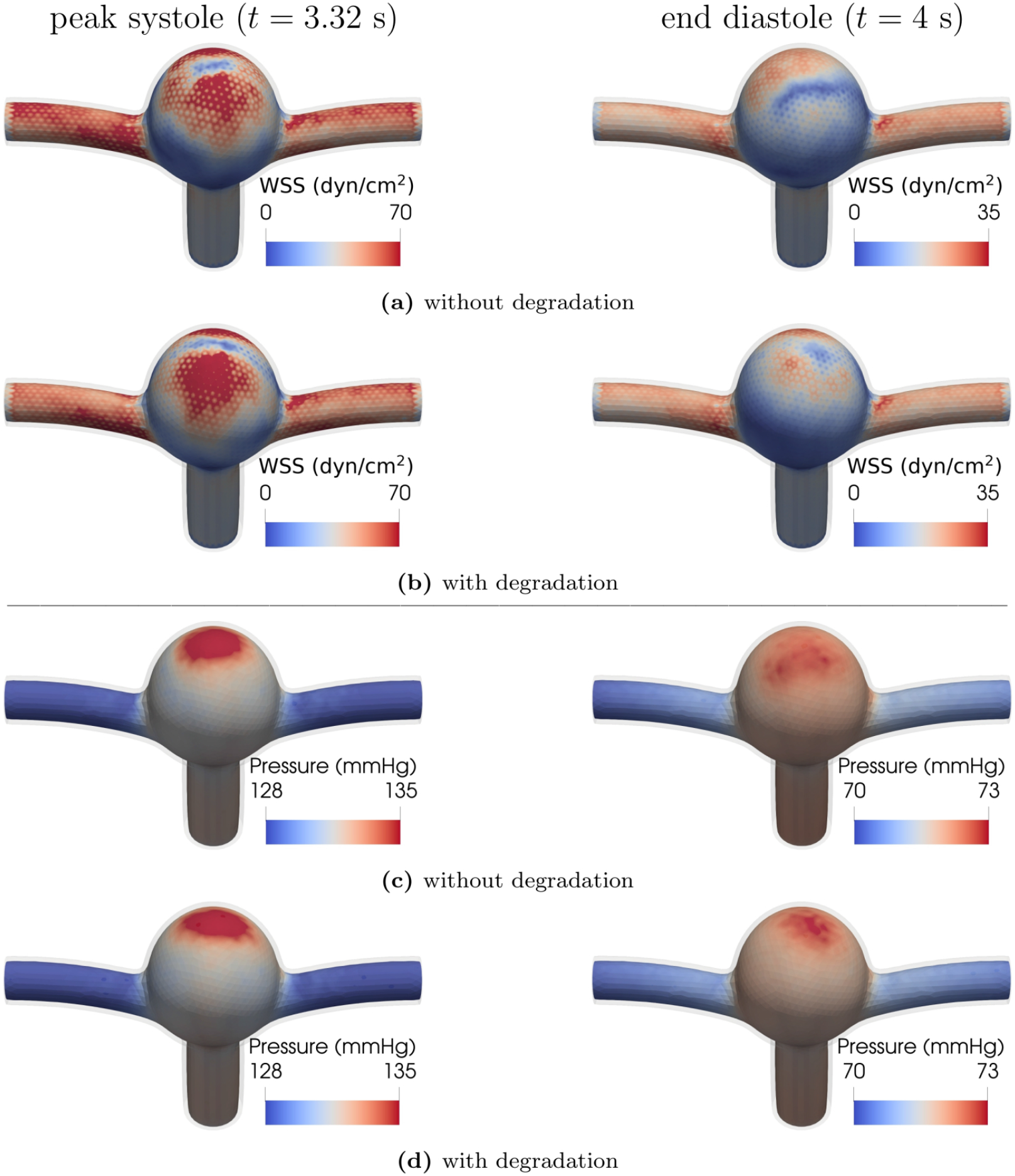
The first two panels show the magnitude of WSS field at (left) peak systole and (right) end diastole (a) without and (b) with tissue degradation. The third and fourth panels present the near-wall pressure field at peak systole and end diastole (c) without and (d) with tissue degradation. Note that the color scales stand for different data ranges in the cases of peak systole and end diastole. The duration of cardiac cycle is *T* = 0.8 s. 1 dyn/cm^2^ = 0.1 Pa.

To highlight the effects of tissue degradation in a more telling way, we use Eq. 8 and determine the *percentage changes* due to degradation in the above-mentioned quantities of interest. Using this approach, the increase or decrease of a quantity upon tissue-degradation shows itself in a positive or negative value, and will be made visible by red or blue color, respectively.

As illustrated in Fig. 4, this analysis reveals that tissue degeneration leads to a reduction of von Mises stress and an increase in strain at the same time. This is direct evidence for a substantial decrease of tissue stiffness due to degradation in the present material model. Thus, the current approach is able to account for *stress softening*, a phenomenon commonly observed in biological tissues.

**Fig 4:**
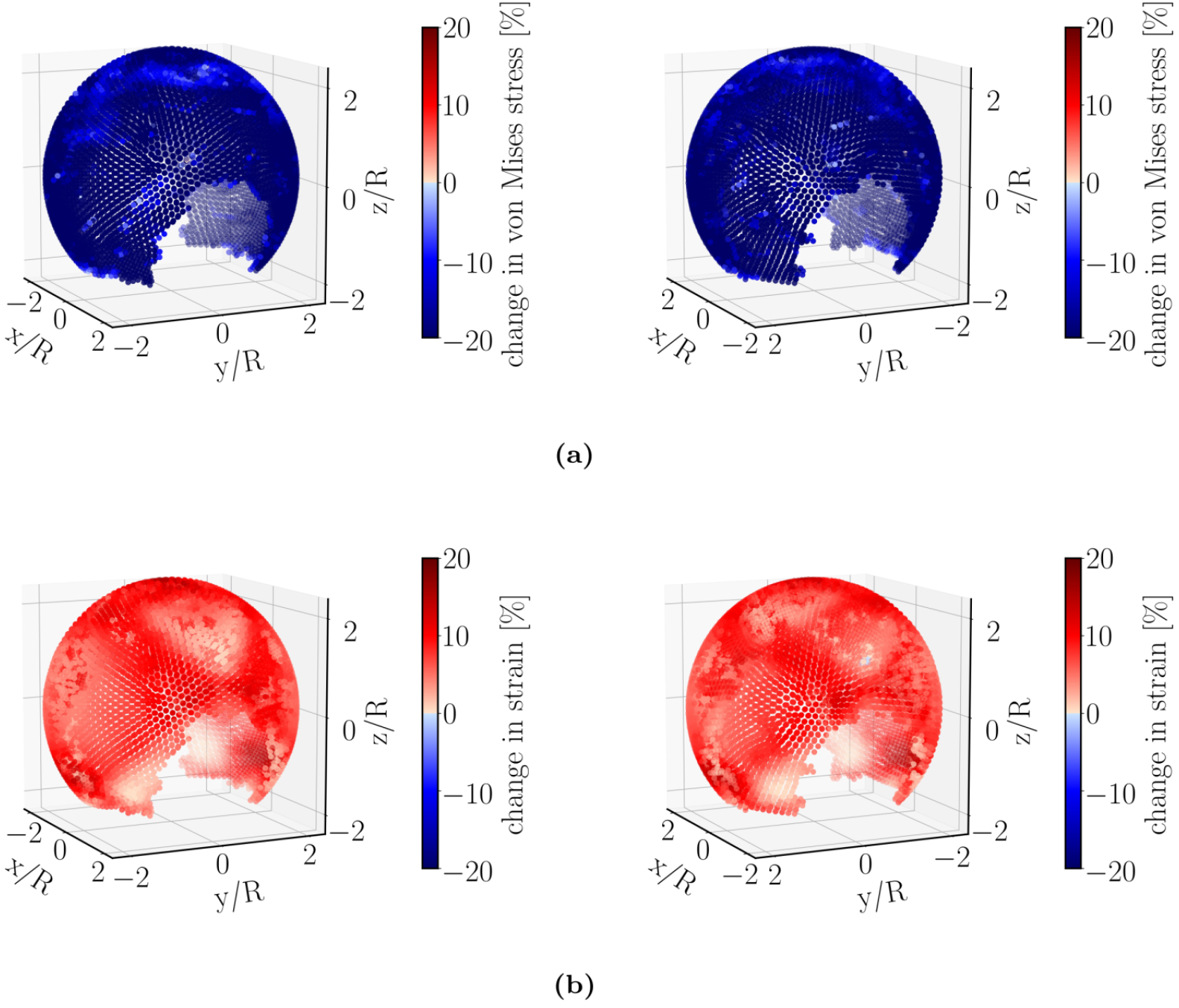
Relative percentage change (Eq. 8) of (a) von Mises stress and (b) strain magnitude of the aneurysmal wall between non-degraded and degraded cases. All quantities are first averaged over a full cardiac cycle (similar to Eq. 7) and then inserted into Eq. 8 to obtain the relative change due to degradation. It is seen from these plots that including degradation leads to an increase in strain and a decrease in von Mises stress. The right panels show the same simulated data as in the left ones but after rotating the aneurysm dome by an angle of *π* around the polar axis.

Moreover, degradation effects seem to be quite heterogeneous. This heterogeneity is strongly enhanced in the case of TAWSS (Fig. 5(a)).

**Fig 5:**
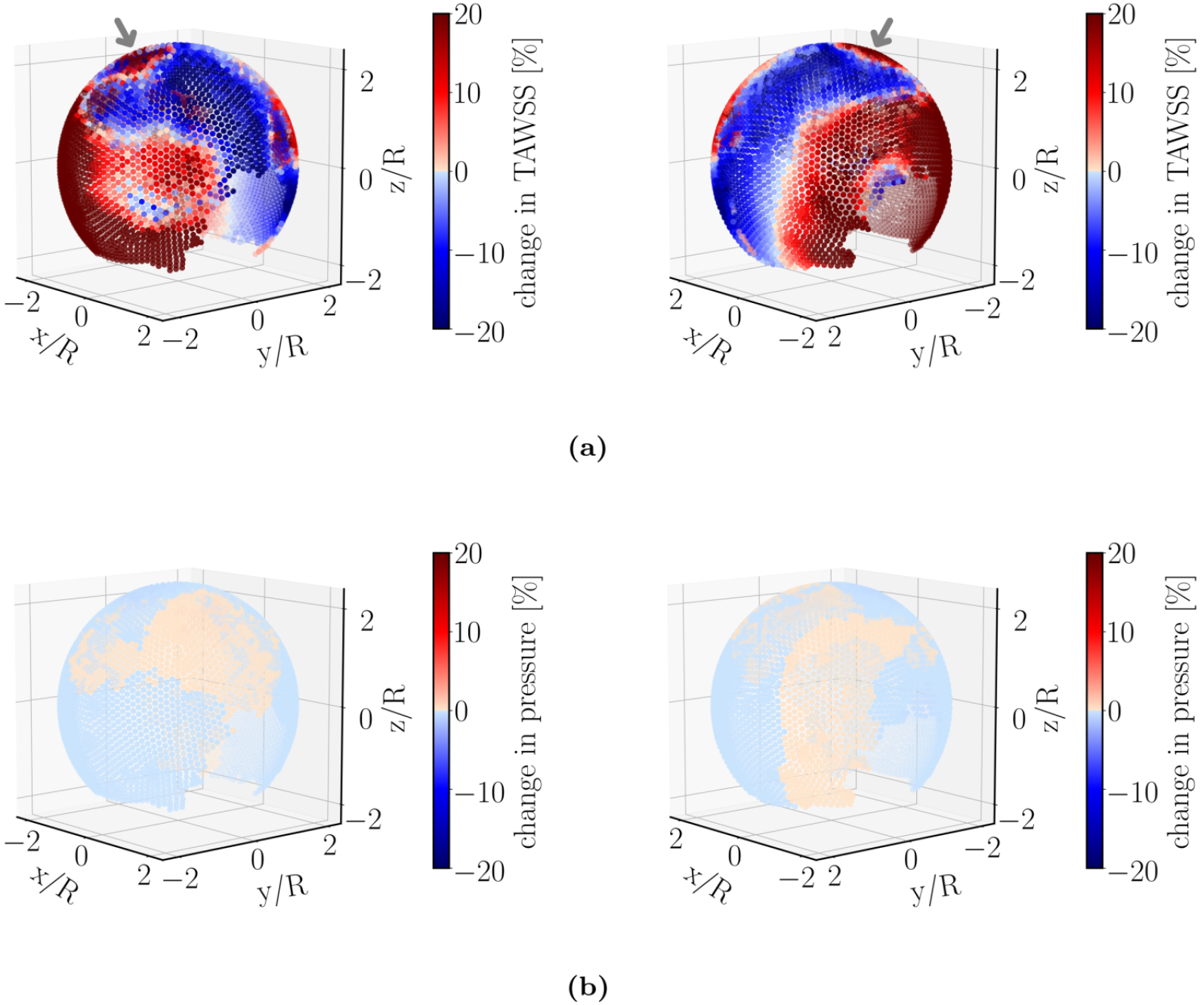
Relative percentage change (Eq. 8) of (a) TAWSS and (b) near-wall pressure. TAWSS shows a significant effect of degradation (roughly 20% variation) and is spatially heterogeneous. Moreover, tissue degradation leads to an increase of TAWSS near the flow-impingement region as indicated by the arrow. Near-wall pressure, however, is rather insensitive to degradation. The right panels show the same simulated data as in the left ones but after rotating the aneurysm dome by an angle of *π* around the polar axis.

In contrast to its rather strong effects on WSS, tissue degradation has a marginal influence on the near-wall pressure. To highlight this issue, we have used the same color scale both for pressure and TAWSS. The small variation in pressure upon degradation thus translates into a pale color in Fig. 5(b). The fluid velocity field inside the bifurcation aneurysm is shown in Fig. 6. With the flow entering the aneurysmal sac, complex flow develops, which impinges on the aneurysmal apex and recirculates inside the dome and leads to the formation of eddies.

**Fig 6:**
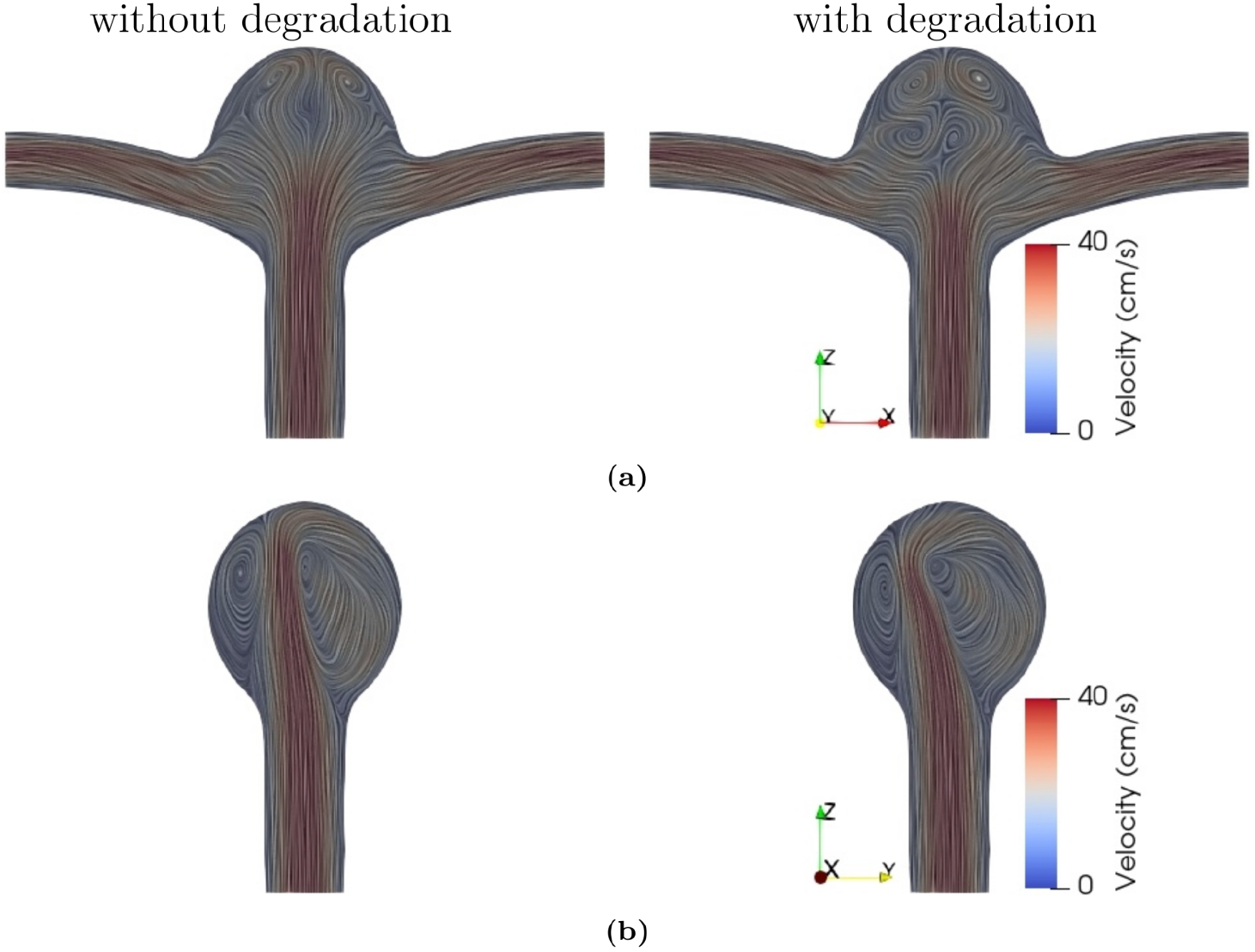
Flow velocity field in the (a) midplane and (b) midway between the two outlets of the bifurcation aneurysm at time instant *t* = 4 s (end diastole), (left) without and (right) with degradation. The duration of cardiac cycle is *T* = 0.8 s. The color scale gives the magnitude of flow velocity. Streamlines serve to visualize the flow pattern.

The impingement of the flow leads to an increase in the local pressure and WSS around the apex of the aneurysm (Fig. 3). This sharp increase in pressure and WSS is of potential clinical interest as it may cause further degradation or even injury to endothelial cells, and thus affect the progression of the aneurysm. Our results indicate that the complexity of the vortical structures is further amplified with tissue degradation (in particular, during diastole as indicated in Fig. 6). We believe that these unsteady variations are primarily responsible for the heterogeneity in WSS.

Furthermore, Fig. 5(a) suggests that tissue-degradation leads to locally large WSS at areas close to the flow impingement site within the aneurysm dome. Irrespective of the details, this feature seems to be robust and persists under different flow conditions, e.g., by varying the blood pressure and/or heart-beat (see Fig. 7).

**Fig 7:**
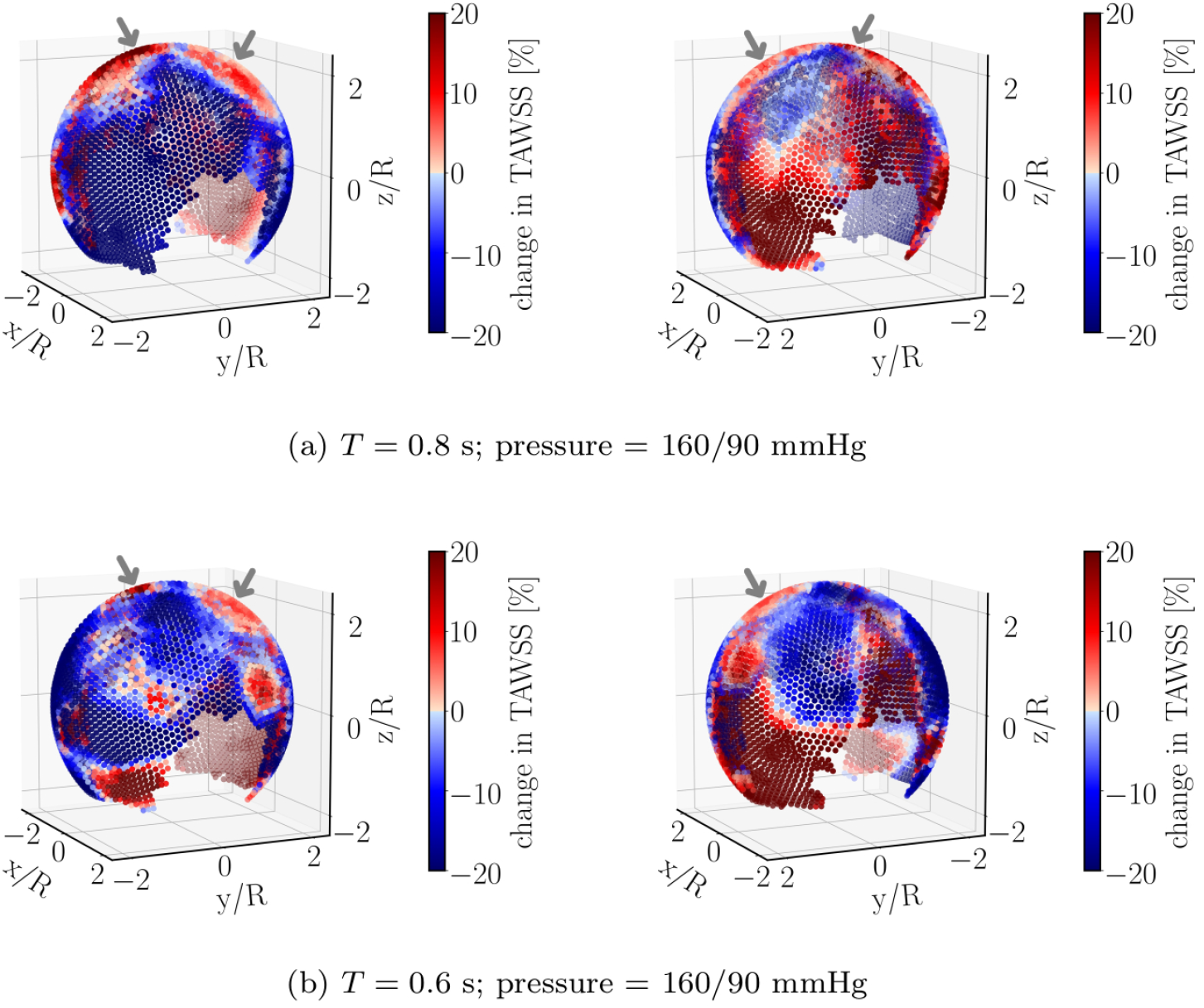
Relative percentage change of TAWSS under different flow conditions. Compared to the data shown in Fig. 5(a), only the range of blood pressure is changed in (a). Panels in (b) differ from (a) in the duration *T* of a single heart-beat. Regions near the flow-impingement are indicated by arrows. The right panels show the same simulated data as in the left ones but after rotating the aneurysm dome by an angle of *π* around the polar axis.

This is an important observation since both high WSS prevalent in the flow-impingement region and low WSS at regions away from it have been associated with the degradation and rupture of aneurysmal walls [7–12, 48].

To examine the role of geometry [14, 37, 49, 50] for the above-described observation, we have conducted simulations of pulsatile flow through a curved tube without aneurysm (Fig. 8) and a different ideal aneurysm (Fig. 9; see also Fig. 1(b)). In the former case, the degradation model is applied to the entire tube. As shown in Fig. 8, changes of TAWSS upon degradation are not noticeable at regions near the flow-impingement site in the absence of vessel abnormalities such as aneurysms. Moreover, we do not observe any hint for a region especially prone to damage.

**Fig 8:**
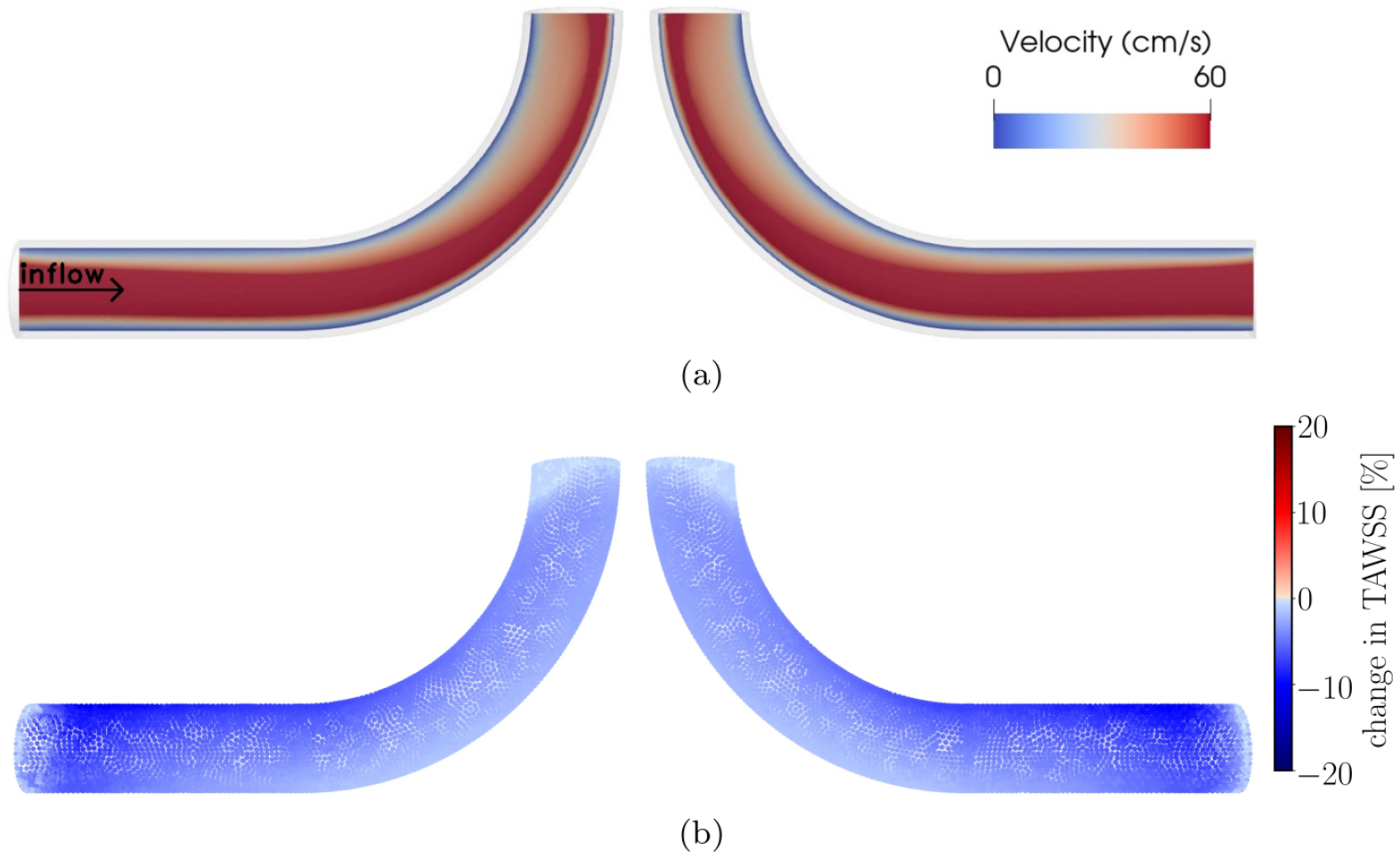
(a) Flow velocity field at peak systole in the presence of degradation within a curved pipe without any aneurysm. (b) Relative percentage change of TAWSS between non-degraded and degraded cases. *T* = 0.8 s. The right panels show the same simulated data as in the left ones but after rotating the geometry by an angle of *π* around the polar axis. Overall, effects on TAWSS associated with degradation and flow impingement vanish in this case.

**Fig 9:**
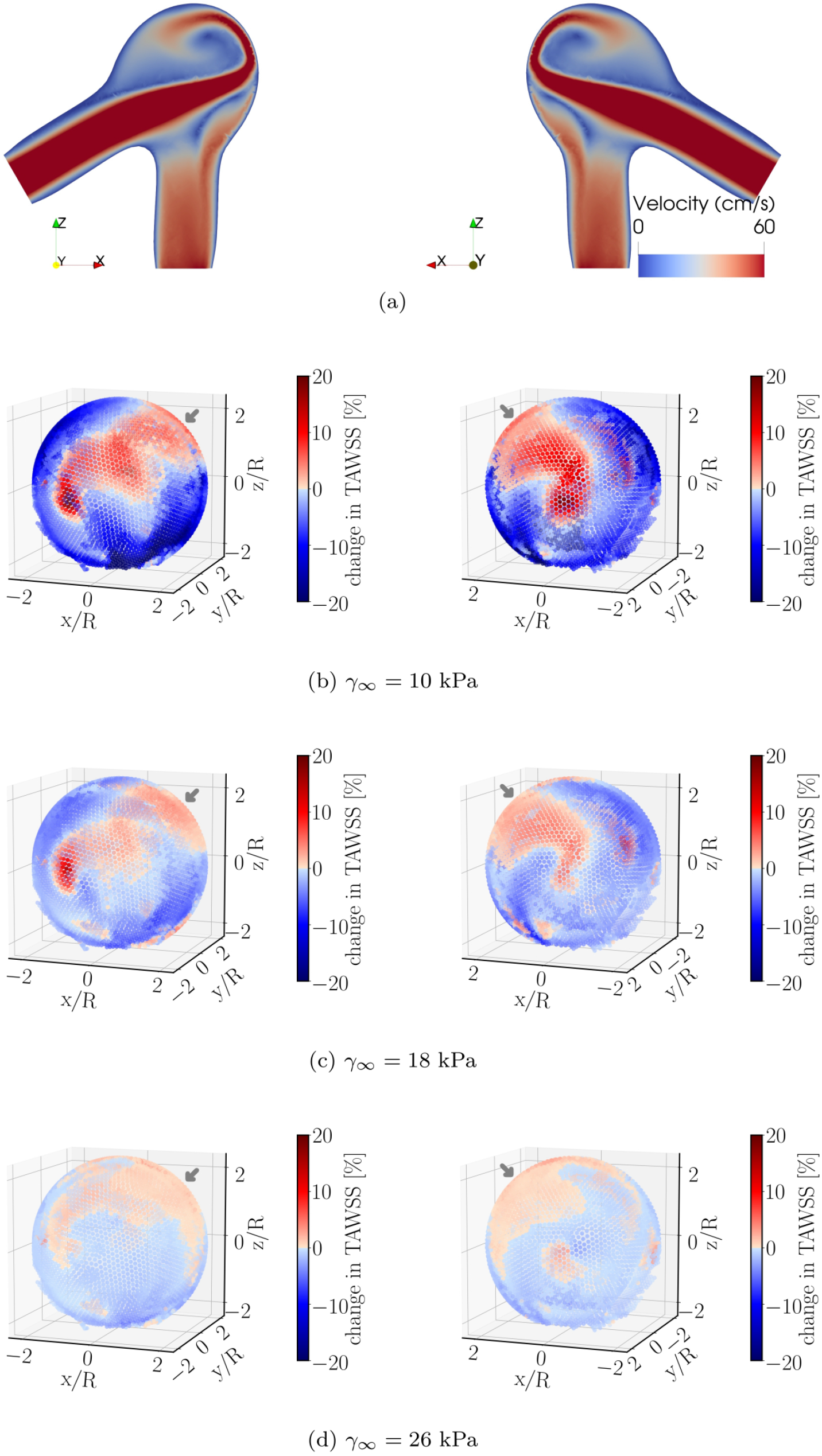
Effect of damage intensity on variations of time-averaged wall shear stress during degradation. (a) Flow velocity field at peak systole in the presence of degradation. The panels (b–d) show the relative percentage change of TAWSS in the aneurysm dome of the geometry (see also Fig. 1(b)). The panels (b), (c) and (d) differ in only the damage intensity via changing the damage parameter *γ*_*∞*_; a small degree of damage is simulated by increasing the value of *γ*_*∞*_ (Fig. 2(b)). *T* = 0.6 s. The region close to the flow-impingement is indicated by the arrow. The right panels show the same simulated data as in the left ones but after rotating the geometry by an angle of *π* around the polar axis.

Finally, we address the effect of degradation intensity on the material’s behavior. For this purpose, we have performed simulations using the same aneurysm geometry but different values of the parameter *γ*_*∞*_. Results obtained for this scenario are depicted in Fig. 9. In qualitative agreement with the data shown in Fig. 5(a) and Fig. 7, local wall shear stress increases upon degradation in a region close to the flow impingement zone. Conversely, far from this zone, degradation leads to a weakening of the stress intensity. All these effects decrease in magnitude with a lower degree of degradation, as characterized by increasing values of the parameter *γ*_*∞*_ (Fig. 2(b)).

### 3.2 Limitations

The numerical approach used in this study deserves several comments. First, the time scale of the degradation process considered in this work was purposely restricted to a few cardiac cycles. This is motivated by the fact that, after the initial degradation stage, no further morphological changes occur with time within the present material model. As visible from Fig. 2(b), the damage saturates after a few (up to a maximum of five) cycles in all the cases investigated. We thus have restricted the computationally expensive FSI simulations to a total of five cardiac cycles. There is no doubt that a comparison between numerical studies and clinical observations is one of the key steps in model development. In the present work, however, we did not find any experimental study which could provide us with the necessary data to enable a complete validation.

Second, it remains a task for future work to account for patient-specific aneurysms and fiber orientations. Indeed, the present degradation model can be improved naturally if experimental and clinical data for specific aneurysms under investigation become available. To the best of our knowledge, however, such data are difficult to be obtained. This is presumably related to the complexity of degradation phenomena and difficulties in accessing the in vivo properties of blood vessel tissue during the degradation process. Direct measurement of the mechanical properties of aneurysmal tissue is a significant undertaking and bears some risk of fatal consequences. Currently, we are not aware of any such clinical experiment. It is noteworthy that we have not used arbitrary, purely academic values for the material parameters and have chosen values in line with the verified parameter adjustments provided in the literature [16]. To face the fact that the particular choice of parameter values does not coincidentally yield the presented results, we analyzed additional scenarios where the parameter values associated with tissue degradation intensity were modified. These calculations show quite similar results and strengthen the conclusions, at least with regard to their qualitative nature (see Fig. 9). The present numerical study thus aims to bridge the gap between the current state of the art and future clinical investigations. We have uncovered an important correlation between flow impingement and a biomechanically important metric, the wall shear stress. This insight may prove useful in designing new experimental studies.

Third, active response and fatigue of the aneurysmal wall are not accounted for in the present material model. It would be interesting to compare a morphological change of an aneurysm after a longtime evolution of degradation. However, computation for a longer time would make sense only if fatigue and/or growth and remodeling are also included in the material model. This remains a challenging task for future studies.

## 4 Conclusion

In this study, we presented a hybrid model which combines a solver for dynamics of blood flow and fluid-structure interactions with a material model for tissue degradation. The coupling scheme was successfully tested and then applied to study how flow under the impact of tissue degradation may influence the mechanical state (which, in turn, may affect fluid flow due to changes in arterial wall geometry and deformability) in different geometries.

The results obtained within this work revealed a strong heterogeneity in the effects arising from degradation on WSS. Importantly, upon degradation, we found increased WSS at places close to the flow-impingement site and decreased WSS beyond the impingement region. This observation is found to be robust and is underlined by simulations of different blood pressures and heart rates. We also present simulated data for a different geometry, confirming this behavior. The set of data provided in this manuscript is thus manifold and supports the proposed connection between effects of degradation and flow impingement on WSS consistently.

Hence, flow impingement seems to be a relevant aspect for a local increase in WSS due to tissue degradation under variable hemodynamic conditions. It would be very interesting to explore possible physiological consequences of this finding for aneurysm’s pathological changes.

## Acknowledgments

This work was performed with support from the IMPRS-SurMat program and by the German Research Foundation (DFG) within SFB-TRR 287/1

